# Cortical reinstatement of causally related events sparks narrative insights by updating neural representation patterns

**DOI:** 10.1101/2025.03.12.642853

**Authors:** Hayoung Song, Jin Ke, Rhea Madhogarhia, Yuan Chang Leong, Monica D. Rosenberg

**Affiliations:** Department of Psychology, University of Chicago, Chicago IL, USA; Department of Neuroscience, Washington University School of Medicine, St Louis MO, USA; Department of Psychology, Yale University, New Haven CT, USA; Neuroscience Institute, University of Chicago, Chicago IL, USA; Institute for Mind and Biology, University of Chicago, Chicago IL, USA

**Author notes:** Correspondence: HS, MDR.

**Keywords:** Narrative comprehension, Insight, Causal inference, Memory retrieval, FMRI

## Abstract

We make sense of everyday events by reasoning about their underlying causes. When we connect causal links between events separated in time, we often experience a sudden feeling of “aha!”, or a moment of insight. What cognitive and neural processes underlie these moments of understanding? We hypothesized that narrative insight accompanies retrieving causally related past events in memory and updating the current event representation. To test this, we designed an fMRI study in which participants watched a TV episode that was cut into multiple events and presented in temporally scrambled orders. Participants pressed an “aha” button whenever they understood something new and verbally explained why they pressed in those moments at the end of each run. Supporting our prediction, more than 40% of insights included the retrieval of past events that were causally related to the current event. Neural patterns representing causally related past events were reinstated in cortical areas. This neural reinstatement drove sudden shifts in cortical representation patterns ∼2 s prior to aha button presses, reflecting an update in situational representation at moments of insight. Moreover, distributed areas in the brain represented causally related events with similar neural patterns, beyond their shared semantic or perceptual features. Together, the study suggests that we comprehend events by reinstating causally related past events via shared neural patterns, followed by updating neural patterns at moments of insight.

## Introduction

We understand everyday events by reasoning about their underlying causes. This process involves not only tracing causal chains of consecutive events but also connecting causes across events separated in time. While causal connections can emerge gradually as events unfold, we often experience a sudden “aha” moment when linking distant events. These moments of insight occur when remembering critical past events. For example, imagine you’re stuck in unusually heavy traffic and can’t figure out why. When you suddenly recall hearing news about a big event happening downtown today, you will experience an “aha” moment and understand the congestion.

In cognitive psychology, insight has been most widely studied in problem-solving (Aru et al., 2023; Becker and Cabeza, 2025), such as when participants solve the nine-dot problem by connecting all dots with four straight, continuous lines (Öllinger et al., 2014). Insight has been defined as a sudden conscious change in a person’s representation of a stimulus, situation, event, or problem (Kaplan and Simon, 1990). This notion was supported by a recent fMRI study that demonstrated changes in neural activity patterns at moments of insight (Becker et al., 2024). Shifts in visual cortical patterns were found when participants suddenly identified difficult-to-recognize Mooney images, which scaled with their reported degrees of insight. In this context, insight was considered as an all-or-none, step-function experience, in contrast to analytic problem-solving which occurs deliberately and incrementally (Kounios and Beeman, 2014).

However, we experience insight not only when we break an impasse to a difficult problem but also when comprehending everyday events and situations. Unlike insight during problem-solving, narrative insight can be achieved through incremental knowledge gain rather than an all-or-none function and often involves reflecting back to past events. To investigate narrative insight, Milivojevic and colleagues (2015) presented participants with clips from *The Sims*, which were either part of coherent narratives or were unrelated to one another. Once participants discovered which clips formed coherent narratives, multivoxel patterns in the hippocampus and medial prefrontal cortex changed, such that neural patterns were more similar for linked versus unlinked events. This finding suggests that narrative insights are characterized not only by neural pattern shifts but also by the reorganization of neural patterns based on memories of related past events. Song and colleagues (2021) conducted a study that directly asked participants to report moments of insight by pressing an “aha” button during narratives. To elicit multiple insights, they segmented movies into discrete events and presented them in a scrambled sequence. The study found that participants experienced insights when an event was highly causally related to past events (although no behavioral measure specific to memory retrieval was collected). This fits with our phenomenological intuition that narrative insight involves linking incoming events with the related past events in memory. However, these studies raise further questions: What specific relationships do these past events have with the current event? How and when do we retrieve related past events during comprehension? More broadly, what neural processes underlie these memory-based insights?

To search for answers, we turned to narrative comprehension literature which has been an active area of study in the late 20th century. To examine the ongoing comprehension process, researchers had participants read narrative texts and instructed them to verbally describe their understanding after reading each sentence or primed them with target words or sentences that appeared earlier in the text to test their recognition (Suh and Trabasso, 1993). Findings showed that participants were more likely to verbally recall and react faster to past probes that were causally related to the current event, which implies an ongoing causal inference process during comprehension. Empirical evidence led to a widely supported theory positing that comprehension is achieved by reinstating causally related past events in memory and constantly updating causal links between events to form a coherent representation of the situation (Trabasso and van den Broek, 1985; Graesser et al., 1994; Zwaan et al., 1995; Zacks and Tversky, 2001; Chen and Bornstein, 2024). This was corroborated by a recent fMRI study by Chang and colleagues (2021). In their study, participants listened to a narrative that alternated between two stories until connections between them were revealed in the latter part of the story. Neural patterns for the related past events were reinstated at these moments of the reveal, which was correlated with how well participants integrated the two stories. Although causal relationships were not directly examined, the work suggests that neural patterns for related past events are reinstated during narrative comprehension.

Linking theories on insight during problem-solving and causal inference during narrative comprehension, we propose three hypotheses about the underlying neural processes. First, we hypothesize that relevant past events are retrieved at moments of insight, reflected as the reinstatement of neural patterns representing these events. Second, we hypothesize that this retrieval is specifically driven by causal relationships, beyond semantic or perceptual similarities between events. Third, we hypothesize that situational representations are updated at moments of insight, marked by sudden shifts in neural patterns. Using an fMRI experiment to test these hypotheses, this study aims to understand how people comprehend ongoing events by gaining insights into their causal connections.

## Results

### Comprehending a scrambled episode involves multiple moments of insight

We collected fMRI data as human participants (N=36; fMRI data of N=33 analyzed) watched a temporally scrambled episode of a TV show, *This is Us* (41m 40s). The episode is an omnibus-style narrative where the stories of four characters—Jack, Randall, Kate, and Kevin, each faced with their own life challenges— unfold in parallel in interleaved order. The four characters’ events play out independently except for the biological twins Kate and Kevin who sometimes appear in the same events. A major twist at the end reveals that Randall was born on the same day as Kate and Kevin and he was adopted, making the three of them triplets. Jack is their father, with his story taking place 36 years in the past. This sudden reveal elicits a big “aha” moment.

We wanted to make the episode difficult to understand so that participants had to actively comprehend the narrative throughout the entire episode while experiencing multiple “aha” moments. To do this, we segmented the episode into 48 events (each 52 ± 21 sec) and scrambled their temporal sequence. In effect, participants watched a sequence of events that jumped back and forth in time, requiring them to connect relational links between events. The 48 events were grouped into 10 blocks in scrambled orders, such that 5 events were assigned to each of the 9 blocks. The 10th block included the last 3 events of the original episode that contained the big reveal. While fixing the order of events within each block, we scrambled the block order to create three sets of stimuli in different block orders. Block 10 was always placed at run 7 for all scrambled-order groups, and blocks 1 to 9 were positioned in the first one-third (run 1-3), middle one-third (run 4-6), and last one-third (run 8-10) of run orders for the three groups (**Figure 1A**). This allowed us to compare behavioral responses and brain activities across scrambled-order groups as participants perceived the same audiovisual stimuli with different knowledge and contexts about the narrative.

**Figure 1.**
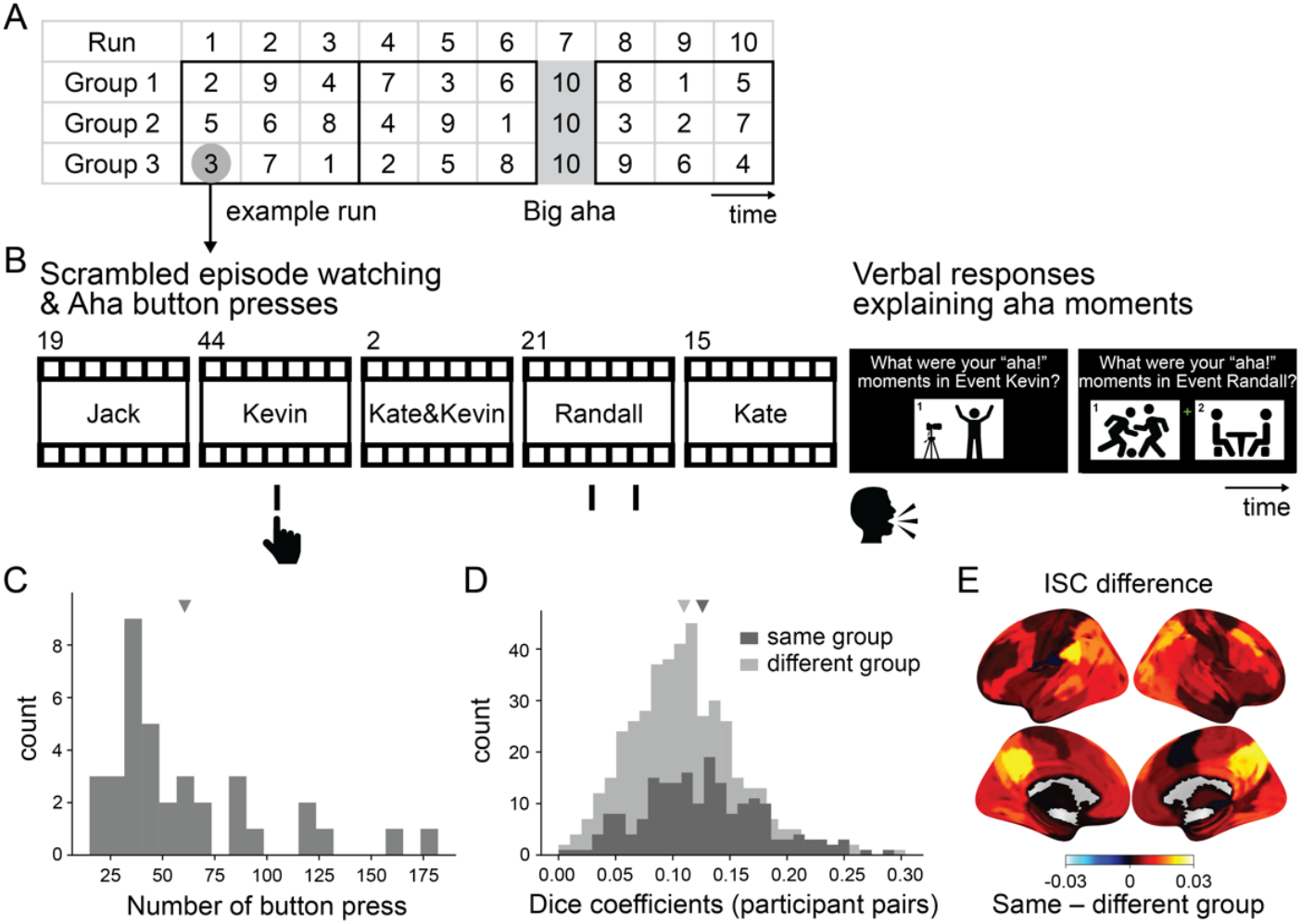
Experimental design and behavioral results. **(A)** Block order for the three stimulus groups. The episode was segmented into 48 events and assigned to one of the 10 blocks. The numbers denote block indices. Blocks 1 through 9 include 5 events in scrambled order, and block 10 includes the last 3 consecutive events of the episode which contain the big reveal. Each block is assigned to either the first, middle, or last one-third of the runs in the three scrambled-order groups (solid contour). **(B)** Experimental design of one example fMRI run. Participants pressed the aha button as they watched scrambled-ordered events from the episode. For each block, an event from Jack, Kate, Kevin, and Randall stories was selected respectively, with one additional event from either Jack event, Randall event, or Kate & Kevin event. The numbers on top of the events denote event indices. Then, participants were presented with screenshots of all the moments they had pressed the aha button and were asked to verbally explain why they pressed in these moments. The screenshots of aha button presses that corresponded to each event were presented together. The schematic figure presents one screenshot for a Kevin event and two screenshots for a Randall event, matching the number of button presses made in respective events. **(C)** Histogram of the aha button presses of 36 fMRI participants. The triangle indicates the mean number of button presses. **(D)** Synchrony in the aha button press moments for all participant pairs. The dark gray histogram indicates the dice coefficients of participant pairs that belong to the same group, whereas the light gray histogram indicates the dice coefficients of participant pairs that belong to different groups. The triangles indicate the mean dice coefficients for respective histograms. **(E)** The difference in mean intersubject correlation (ISC) values across participant pairs that belong to the same scrambled-order group versus different groups. ISC was calculated for 100 cortical and 16 subcortical parcels, respectively.

Thirty-six participants were randomly assigned to one of the three scrambled-order groups (12 participants per group). Participants were instructed to press an “aha” button whenever they understood something new about the show’s events and characters. After watching a set of events in a run, participants were shown screenshots taken whenever they pressed the aha button and were asked to verbally explain why they pressed the button at these moments (**Figure 1B**). The motivation was to know the exact moments when, and the reasons why, participants experienced insight.

The frequency of aha button presses varied across participants. On average, participants pressed the button roughly once every 41s, with a median of one press every 58s, ranging from one press per 14s to 2m 47s (**Figure 1C**). Calculating dice coefficients of participant pairs’ button press moments showed that participants pressed the aha button at similar times, significantly higher than chance (*z* = 35.249, *p* < .001; **Figure 1D**). The aha button presses were more synchronized among participants within the same scrambled-order group compared to different groups (Wilcoxon rank-sum test, *z* = 3.617, *p* = .0003; **Figure 1D**), which illustrates that insight is not only driven by the stimulus but also modulated by the context and the amount of prior knowledge. This behavioral result was complemented by the fMRI result, where a majority of parcels exhibited positive intersubject correlation (ISC) between pairs of participants, and higher within-group ISC compared to across-group ISC (98 out of 100 cortical parcels and 11 out of 16 subcortical parcels based on ISC difference). Together, results show that participants experienced multiple insights at similar moments in time when comprehending the scrambled episode.

### Causally related past events are retrieved at insight moments

Why did participants press the aha button in those moments? To understand this, we analyzed participants’ verbal explanations of their aha button press moments. Based on narrative comprehension theory, we hypothesized that participants retrieved past events that were causally related to the current event (Trabasso and van den Broek, 1985; Graesser et al., 1994; Zwaan et al., 1995).

For a data-driven, qualitative understanding of insight moments, three of the authors labeled each verbal explanation of insight moments into a set of categories (**Supplementary Table S1**). The categories were not defined *a priori* but were qualitatively derived by the first author to maximally cover the spectrum of participants’ verbal responses. Each response could be associated with multiple categories; for example, a participant may have had an aha moment about what is going on in the current event (current event understanding) based on their memory of a past event (memory retrieval) while speculating about what may happen next (inference). We did not include the 7th run in the analysis because most of the responses were about their “big aha”, which was hard to disentangle into categories. Coding of verbal responses found that 41.15 ± 16.64 % of the aha explanations included mentions of past events (ranging from 12.00 to 76.79 % across participants), even when this was not an explicit instruction given to participants. The result indicates that many insight moments were achieved by comprehending the current event in relation to past events.

Then *what* past events were retrieved in memory? Were they causally related past events as we hypothesized? To ask this, we created a memory retrieval matrix by coding past events that were retrieved when participants explained their insight moments. If a past event (or events) was mentioned when explaining an aha button press at a current event, +1 was assigned to that pair(s) of events in an event-by-event matrix (interrater reliability among three raters: pairwise Spearman’s *rho* = .803, .596, .640, *p* values < .0001 compared to chance created from shuffled matrices). This memory retrieval matrix was created for each participant, temporally unscrambled to its original sequence, and then averaged across all participants to create a single memory retrieval matrix (**Figure 2A**). This matrix thus represents the degree to which every event pair is related in memory, or how likely an event is to prompt retrieval of, or to be retrieved during, another event. Additionally, we created a causal relationship matrix by rating the causal relationship between every pair of events on a scale of 0 (No causal relationship) to 4 (Very high causal relationship) (interrater reliability among three raters: pairwise *rho* = .701, .754, .818, *p* values < .0001) (Trabasso et al., 1989; Antony et al., 2024) (**Figure 2B**; **Supplementary Table S2**). When comparing the memory retrieval matrix with the causal relationship matrix, we found a significantly high positive correlation (Spearman’s *rho* = .715, *p* < .0001 compared to chance created from shuffled matrices), meaning memory retrieval followed the causal event structure.

**Figure 2.**
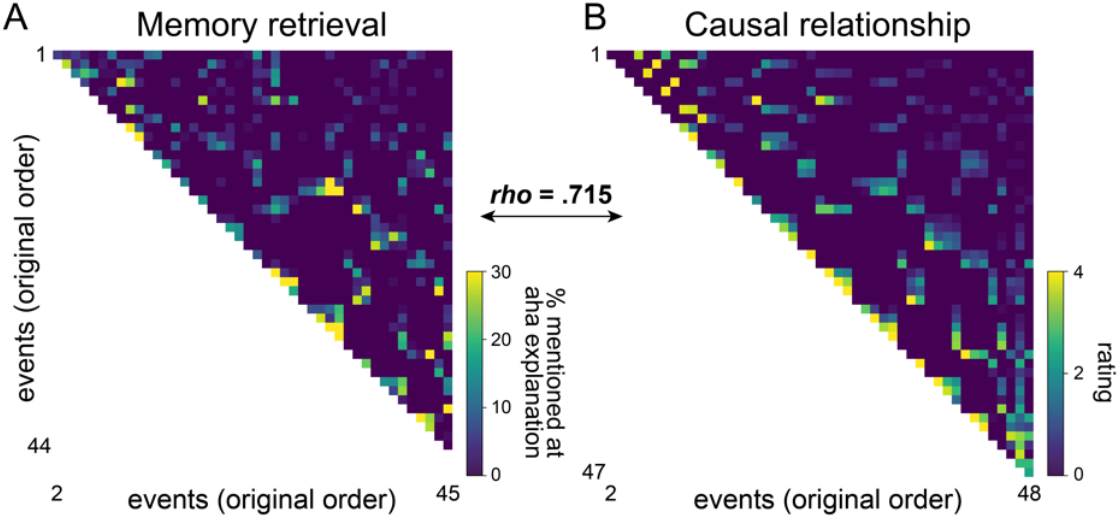
Relationship between past event retrieval at insight moments and event causality. **(A)** Memory retrieval matrix indicates the likelihood of an event retrieving, or being retrieved by, another event when participants explained their aha button presses. The events were unscrambled in order, and memory retrieval matrices created for 32 participants were averaged. Events 46 to 48 (in 7th run) were not included in analyses. **(B)** The causal relationship matrix indicates the degree of causal relationship rated for every event pair on a scale of 0 to 4.

To consider alternatives other than the causal relationship, we created various narrative feature similarity matrices, including similarities in semantic contents of event annotations, which characters appeared in the event, where the event took place, low-level visual features (hue, saturation, pixel intensity, and motion energy of the video), and low-level audio features (amplitude and pitch of left and right stereos) (**Supplementary Figure S1A**). We also created a time matrix indicating how nearby in time the event pairs are in its original sequence. The causal relationship matrix and all alternative features we considered, with the exception of audio features, were similar to one another (pairwise *rho* values ranging from .216 to .858, FDR-corrected *p* values < .0001) (**Supplementary Figure S1B**). A multiple linear regression analysis found that the causal relationship matrix and the 6 narrative feature matrices together predict memory retrieval matrix, with *R*^2^ = .527, where causal relationship explained the largest variance (*t* = 16.474, *p* < .001; standardized coefficient = .615) amongst all variables (*t* values ≤ 2.517; coefficients ≤ .087). Importantly, when predicting the memory retrieval matrix from the causal relationship matrix alone, the explained variance remained less changed, with *R*^2^ = .519 (model comparison, *F* = 2.664, *p* = .014). On the contrary, when predicting memory retrieval matrix from the 6 narrative feature matrices without the causal relationship matrix, the explained variance showed a pronounced decrease, with *R*^2^ = .396 (*F* = 271.387, *p* = 5.05e-54). This indicates that participants retrieved causally related events in memory at insight moments, which was beyond similarities in semantics, characters, places, and low-level audio and visual features, as well as proximity in time.

### Cortical representation patterns shift at moments of insight

What neural signature characterizes the experience of insight? Motivated by the psychological definition of insight as a sudden conscious change in representation (Kaplan and Simon, 1990) and in accordance with recent neuroscientific findings (Becker et al., 2024), we hypothesized that neural activity patterns would suddenly change at moments of insight. To test this, we used a data-driven hidden Markov model (HMM) to identify moments of sudden shift in voxel activity patterns (Baldassano et al., 2017) (**Figure 3A**). The HMM was applied to multivoxel activity time series in each of the 100 cortical parcels (Schaefer et al., 2018), for each of the 48 events and 33 participants respectively.

**Figure 3.**
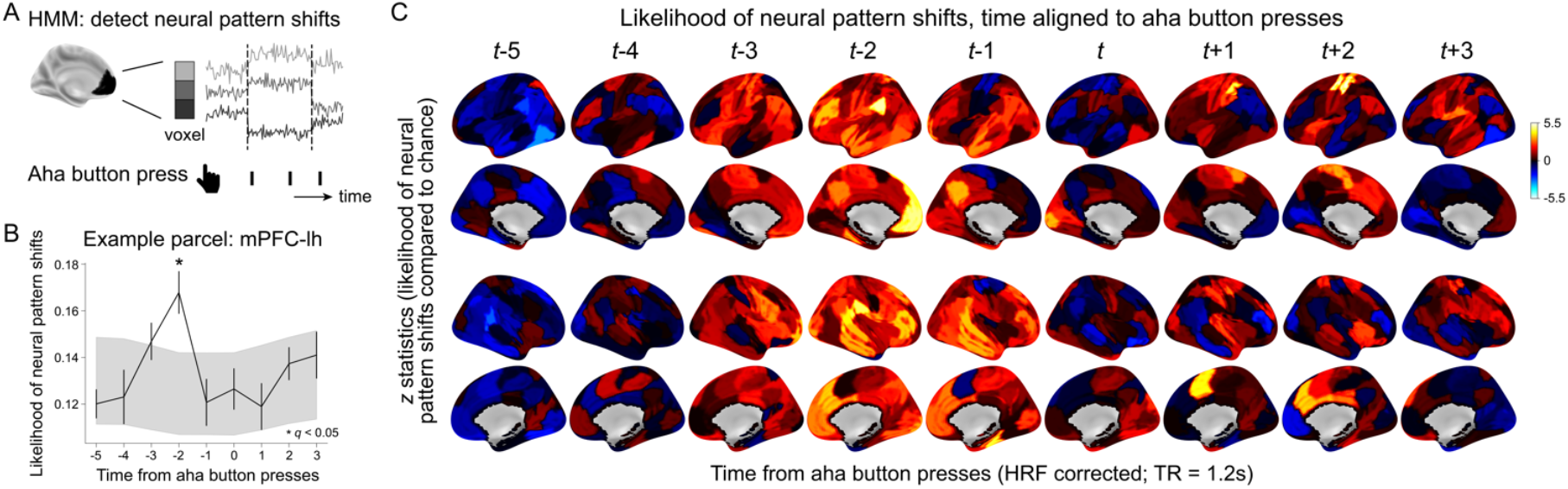
The likelihood of neural pattern shifts near aha button press moments. **(A)** Schematic illustration of the method. For each parcel, the hidden Markov model (HMM) was applied to identify sudden shifts in multivariate BOLD activity patterns. Moments of HMM-detected neural pattern shifts were compared to HRF-shifted aha button presses, based on the hypothesis that insight moments would correspond to changes in neural patterns. **(B)** Likelihood of neural pattern shifts compared to a chance distribution with randomly shuffled aha button presses, visualized in an example parcel, the left medial prefrontal cortex (mPFC). The solid line indicates the likelihood of neural pattern shifts (± standard error of the mean). The gray area indicates the 95% confidence interval of the chance distribution (10,000 iterations). Asterisks indicate time points significantly different from chance (corrected for the number of parcels and time steps, FDR-*p* < 0.05). **(C)** The same result as **B** but in 100 cortical parcels. Colors indicate z statistics computed in comparison to the chance. **Supplementary Figure S2** shows a thresholded *z*-statistics map at FDR-*p* < 0.05.

We hypothesized that participants’ aha button presses would be aligned with moments of neural pattern shift detected by the HMM. To test this, we time-aligned the HMM-detected shifts to a participant’s every aha button press moment (−5 to 3 TRs from the aha button response times, after shifting 4 TRs [4.8 s] to account for hemodynamic response delay [HRF]), which were averaged across all button press moments. The likelihood of neural pattern shift from -5 to 3 TRs from aha button presses were compared to a chance distribution where the button press moments were randomly shuffled (**Figure 3B**).

**Figure 3C** shows *z* statistics, indicating the likelihood of representation pattern shifts compared to the chance distribution, of 100 cortical parcels, at -5 to 3 TRs from the HRF-shifted aha button presses. An overall increase in the likelihood of neural pattern shifts was observed across distributed areas in the cortex at ∼2 s prior to aha button presses. Specifically, 19 out of 100 cortical parcels indicated a sudden shift in representation patterns at 2 TRs (=2.4 s) prior to button presses (see **Supplementary Table S3** for a list of these 19 regions), and 8 out of 100 cortical parcels at 1 TR (=1.2 s) prior to button presses (see **Supplementary Figure S2** for a thresholded map, FDR-*p* < .05). All of these significant parcels showed positive *z* statistics. Regions indicated to be significant bilaterally were the dorsolateral prefrontal cortex, parietal operculum, parahippocampal cortex, and the temporal lobe. The number of significant parcels detected were ≤ 4 out of 100 parcels for the rest of the time points tested. The results indicate that representation patterns of the distributed cortical areas shifted at roughly 2 s prior to participants’ aha button press moments. The result suggests that the change in cortical representation patterns characterizes the experience of insight, reflecting an update of the situational representation (for the relationship between insights and perceived event boundaries, see **Supplementary Text S1**).

### Neural reinstatement of causally related past events drives neural pattern shifts at insight moments

What drives neural pattern shifts at moments of insight? Based on behavioral results, we asked whether the reinstatement of causally related past events is related to the neural pattern shift. To calculate neural reinstatement, we summed neural patterns of past events, weighted by their causal relationships with the current event. This integrated neural pattern representing causally related past events was correlated with the neural pattern near moments of aha button presses, representing the degree of neural reinstatement (see ***Methods***). We focused our analysis on the 19 cortical regions that significantly shifted neural patterns 2 TRs prior to aha button presses.

**Figure 4A** visualizes neural reinstatement time courses aligned to moments of aha button presses. The aha button presses were grouped into 4 categories based on whether i) a neural pattern shift was detected 2 TRs prior to the aha button press, and ii) whether a participant mentioned past events when verbally explaining their insight for this button press. A neural reinstatement effect was observed from approximately 7 (=8.4 s) to 3 TRs (=3.6 s) before the aha button presses. The effect was pronounced when a neural pattern shift occurred 2 TRs before the button press and when the press was associated with behavioral retrieval of causally related past events (solid green line in **Figure 4A**). When neither of these occurred, the average neural reinstatement effect was near zero (dashed orange line in **Figure 4A**).

**Figure 4.**
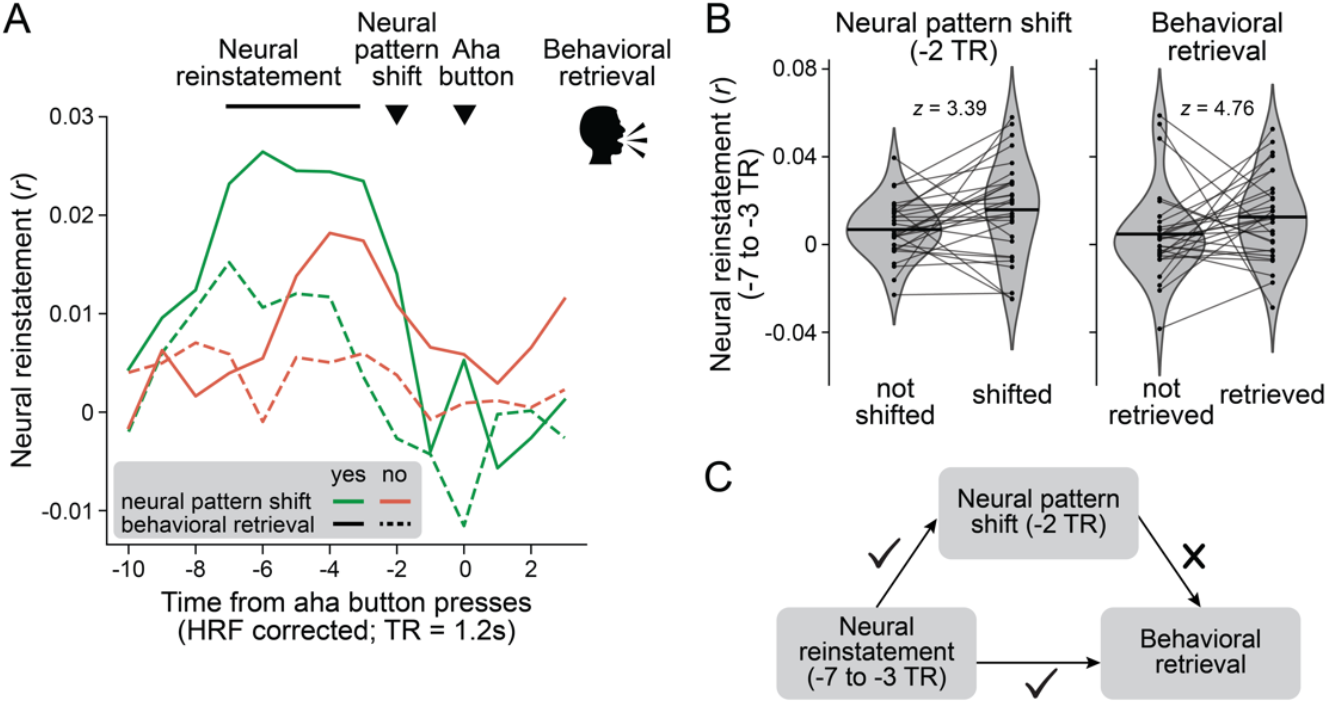
Relationship between neural reinstatement of causally related past events, neural pattern shifts, and behavioral retrieval of causally related past events. **(A)** Neural reinstatement time aligned to HRF-shifted aha button presses. The aha button presses were categorized based on whether neural shifts occurred 2 TRs prior to the button presses and whether past events were behaviorally retrieved during verbal response phases (category indicated by colors and styles of the lines). Lines indicate the average across 19 regions and 29 participants. **(B)** Mean neural reinstatement at -7 to -3 TRs from the aha button presses, with respect to whether the neural patterns shifted vs. not at -2 TR (*left*), and whether past events were retrieved vs. not when explaining the corresponding button presses (*right*). Dots indicate the average neural reinstatement of 19 regions for each of the 29 participants. Lines within the violin plot show the means of the distributions. Statistics were conducted using mixed effects models to account for the effects of 19 regions and the random effects of participants. **(C)** Schematics of the results. The neural reinstatement significantly predicts both the neural pattern shift and behavioral retrieval, whereas the neural pattern shift does not reliably predict behavioral retrieval.

To test this statistically, we asked whether the degree of neural reinstatement influences the likelihood of neural pattern shift (**Figure 4B**). We used a logistic generalized linear mixed effect model to predict neural pattern shifts at -2 TR, given the average neural reinstatement effect at -7 to -3 TRs from the aha button presses. Neural pattern shifts were more likely to be detected when the neural reinstatement was high (χ^2^(1) = 11.475, *p* = .001, *z* = 3.388), indicating that the neural reinstatement of causally related past events influenced updates in neural patterns.

We further asked if neural reinstatement, neural pattern shifts, and their interaction predict behavioral retrieval of causally related past events. Neural reinstatement predicted behavioral retrieval (χ^2^(1) = 15.268, *p* = 9.3e-05, *z* = 3.905), corroborating neural reinstatement as a measure of memory retrieval. We also found a main effect of neural pattern shift at -2 TR predicting behavioral retrieval in the negative direction (χ^2^(1) = 9.075, *p* = .003, *z* = -3.016), although this effect was gone when we included neural pattern shifts that occurred at -1 TR (χ^2^(1) = .676, *p* = .411, *z* = -0.822) which refrained us from making further interpretations. No interaction was found between neural reinstatement and neural pattern shift in predicting behavioral retrieval (χ^2^(1) = 1.277, *p* = .259, *z* = 1.130).

Together, we found two distinct neural signatures involved in scrambled narrative comprehension. Neural reinstatement reflects memory retrieval of causally related past events and neural pattern shift reflects updates in situational representation at moments of insight. Neural reinstatement occurred prior to neural pattern shifts and was a significant predictor for both neural pattern shifts and behavioral retrieval. However, the neural pattern shift itself—nor its interaction with neural reinstatement—did not exhibit a meaningful relationship with behavioral retrieval. This corroborates our behavioral finding: Comprehension based on retrieval of causally related past events is one among many different reasons for insight experience. Reinstatement of causally related past events drives insight, but having experienced insight does not necessarily mean that past events are retrieved in memory (**Figure 4C**).

### The brain represents causally related events with similar neural patterns

Neural patterns representing causally related past events were reinstated before moments of insight, as indicated by an increased positive correlation between past and present neural patterns. This suggests that causal relationships between events are encoded through shared representation patterns in the brain. We investigated i) whether this representational similarity extends beyond nearby moments of insight, and ii) whether the effect is specific to causal relationships rather than other shared features between events. We employed representation pattern similarity analysis (Kriegeskorte et al., 2008) to ask if causally related events were represented by similar BOLD activity patterns.

The BOLD activity time series were extracted from all voxels in each of the 100 cortical (Schaefer et al., 2018) and 16 subcortical (Tian et al., 2020) parcels as participants watched 48 events in scrambled orders. The BOLD time series were averaged across time for 48 events respectively, resulting in a (number of voxels) x (48 events) representation pattern matrix. A Pearson’s correlation coefficient was computed to estimate the event-by-event BOLD pattern similarity matrix (**Figure 5A**). To avoid spurious correlations, we only computed similarities between events that belonged to different fMRI runs (Mumford et al., 2014). The BOLD pattern similarity matrix was correlated with the causal relationship matrix (in the order that scrambled events were presented) using Spearman’s rank correlation. Compared to chance distributions created by shuffling the BOLD pattern similarity matrix, all 100 cortical parcels and 9 out of 16 subcortical parcels exhibited significantly positive correlation above chance (FDR-*p* < 0.05), meaning causally related events were represented by similar neural patterns across the whole brain.

**Figure 5.**
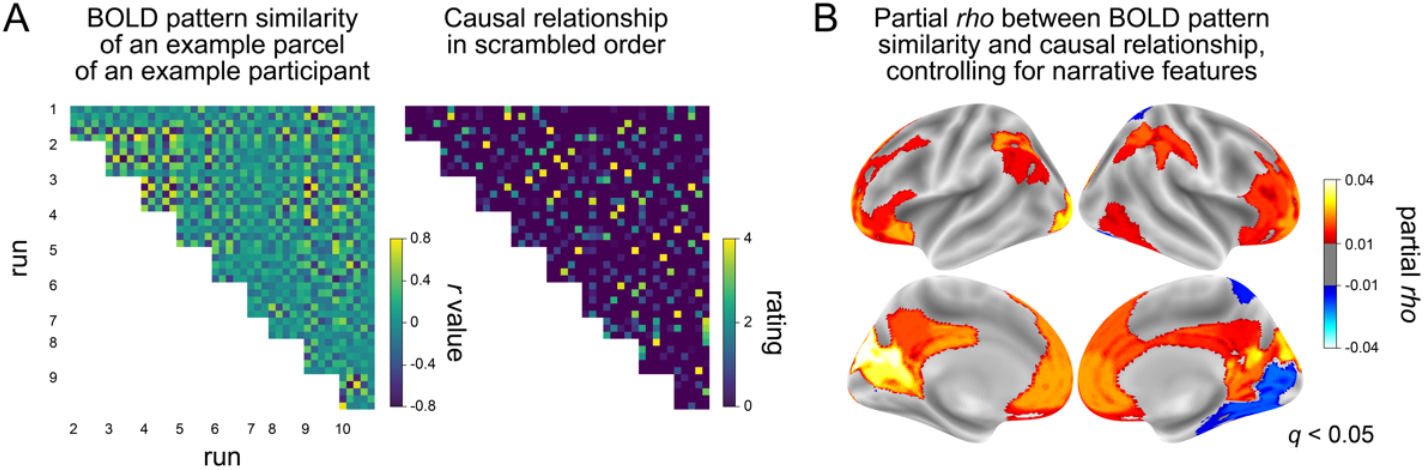
Relationship between neural activity pattern similarity and causal event structure. **(A)** (*Left*) The event-by-event BOLD activity pattern similarity matrix of an example parcel of an example participant. (*Right*) The same causal relationship matrix as in **Figure 2B** but in scrambled order. **(B)** Parcels that hold a significant correlation between neural pattern similarity and causal relationship, while controlling for various narrative features such as semantic and perceptual similarities (thresholded at FDR-*p* < 0.05). Colors indicate partial *rho* values averaged across participants.

Because causal relationships between events are intrinsically correlated with various narrative feature similarities and temporal proximity, we computed a partial correlation between BOLD pattern similarity and causal relationship, while controlling for narrative features including semantic and perceptual similarities and temporal distance (**Supplementary Figure S1**). Even after this strict control, 40 out of 116 parcels significantly represented causally related events with similar neural patterns, among which 36 out of 40 parcels exhibited positive partial *rho* values (FDR-*p* < 0.05) (**Figure 5B**). These regions included the bilateral retrosplenial cortex and the nearby visual areas which exhibited the largest *rho* values, and the bilateral mPFC, ventrolateral and dorsal prefrontal cortex, intraparietal cortex, and precuneus and posterior cingulate cortex. This suggests that wide regions in the brain use representation patterns to construct a causally coherent event structure, beyond semantic and perceptual similarities.

## Discussion

This study investigated how people comprehend narratives by piecing together causal links between events at multiple moments of insight. FMRI participants pressed an aha button while watching a temporally scrambled episode and verbally explained why they pressed in those moments at the end of each run. Participants experienced insights during comprehension, many of which included the retrieval of past events. Consistent with event comprehension theory, our behavioral results showed that causally related past events were reinstated in memory, suggesting an ongoing process of causal inference during comprehension. Aligned with the psychological notion of insight as a representational update, sudden shifts in cortical representation patterns were observed approximately two seconds before the aha button presses. Neural reinstatement of causally related past events—a neural signature of memory retrieval— occurred prior to these neural pattern shifts and predicted their likelihood. Finally, distributed areas in the brain represented causally related events with similar neural patterns, suggesting a way in which we form a causal understanding of narratives. In sum, the study provides neural evidence for how we reinstate memories to connect causal links between events and update situational representations at moments of insight.

In this study, participants indicated their moments of insight while watching a temporally scrambled narrative and provided verbal explanations for each insight at every run. This paradigm allowed us to examine neural processes underlying memory retrieval, causal inference, and insight during narrative comprehension. Behavioral findings showed that more than 40% of participants’ insights included references to past events. The retrieved past events were the events that held strong causal relationships with the current event, with the correlation between memory retrieval and causal relationships exceeding 0.7. While causal relationships between events were intrinsically positively correlated with various narrative features, including semantic and perceptual similarities, causal relationships accounted for the largest variance in memory retrieval. These findings provide direct support for the theory that views comprehension as a process of retrieving causally related past events to update causal links between events (Trabasso and van den Broek, 1985; Graesser et al., 1994; Zwaan et al., 1995; Zacks and Tversky, 2001; Chen and Bornstein, 2024). Insight, as operationalized in this study, can be interpreted as the process of constructing and updating a coherent situational representation by retrieving causally related past events.

Cognitive theories have characterized insight as a sudden change in representation (Kaplan and Simon, 1990). Supporting this notion, we found shifts in neural patterns at approximately 2 seconds prior to aha button presses in distributed areas of the cortex. Drawing on prior studies (Milivojevic et al., 2015; Becker and Cabeza, 2025; Sava-Segal et al., 2025), we propose that these shifts may reflect changes in narrative representation occurring at moments of insight. However, this interpretation requires caution, as neural pattern shifts could also arise from alternative processes, such as motor preparation, motor execution, or decision-making. Furthermore, a prior study that used the same HMM analysis found neural pattern shifts at perceived event boundaries (Baldassano et al., 2017; Geerligs et al., 2021). This is supported by the results of our additional behavioral experiment, which showed a significant correspondence between moments of insight (“aha” button presses) and moments of perceived event boundaries (“event boundary” button presses) (**Supplementary Text S1**). This raises an interesting hypothesis—whether experiencing insight creates internally-driven event boundaries. We encourage future work to further examine the distinction between external and internal event boundaries, and how internal event boundaries relate to the experience of insight.

Participants retrieved causally related past events at moments of insight, an experience that was characterized by shifts in cortical patterns. Based on this, we asked whether the neural patterns of causally related past events are reinstated near moments of insight and whether they have consequences to neural pattern updates. To calculate neural reinstatement, we summed neural patterns of past events after weighing them by their causal relationships with the current event, and then correlated this integrated representation with the neural pattern of the current moment near button presses. This approach was motivated by seminal theories on memory integration (Zeithamova et al., 2012; Milivojevic et al., 2015; Schlichting and Preston, 2017), although different integration mechanisms can be explored. In cortical areas that showed significant neural pattern shifts at 2 TRs prior to aha button presses, we found the neural reinstatement effect at approximately 7 (=8.4 s) to 3 (=3.6 s) TRs prior to the aha button presses. Intriguingly, the neural reinstatement effect was pronounced when neural patterns shifted and when there was behavioral evidence of memory retrieval. The neural reinstatement effect not only predicted neural pattern shifts but also predicted whether participants would mention past events during the verbal response phase. To the best of our knowledge, this is the first study that has demonstrated the relationship between memory retrieval and insight during narratives: reinstatement of causally related past events occurs first, followed by a sudden change in neural patterns which marks moments of insight. Neural reinstatement predicts neural pattern shifts, but neural pattern shifts do not necessarily predict behavioral memory retrieval. This supports our behavioral observation that memory retrieval-based insight is one among various reasons why insight occurs (e.g., insights about current events, characters’ traits and behaviors, and relationships between characters).

Moreover, distributed areas of the brain were found to represent causally related events with similar neural patterns. This remained true even after controlling for narrative features, including similarities in semantic contents, characters, places, low-level visual and audio features, as well as proximity in time. This indicates that the brain encodes causal event structure by representing causally related events with more similar neural patterns and less similar for unrelated events. Future work can examine whether shared neural patterns are driven by transient neural reinstatement of the causally related past events or structured reorganization of neural patterns that are sustained over time.

In sum, this study investigated the cognitive and neural processes underlying narrative insight, guided by memory and ongoing causal inference. Many insights were driven by the retrieval of causally related past events, supported by the neural reinstatement of these past events. This neural reinstatement drove changes in neural patterns at moments of insight which suggest updates to the situational representation. Finally, causally related events were represented by similar neural patterns, suggesting a way in which the brain encodes causal event structure in narratives.

## Acknowledgment

We thank Rachael A. Young and Megan E. Dorn for help with fMRI data collection. We thank Todd B. Parrish for developing an MRI-compatible voice recording device. We thank Emma Tung, Taysha Martinez, Anurima Mummaneni, Jadyn Park, Julia Pruin, Alisa Schutz, and Lucy Tindel for help with behavioral data coding. We thank JeongJun Park and Jungwoo Kim with conceptualization and helpful discussions and comments on the manuscript. The research was supported by the National Science Foundation BCS-2043740 (MDR), Social Sciences Research Center Faculty Seed Grant Program at the University of Chicago (MDR and YCL), and American Psychological Association Dissertation Research Award (HS).

## Data and code availability

Raw and processed fMRI data will be shared in OpenNeuro and behavioral data and codes will be shared in GitHub upon publication.

## CRediT author statement

HS: Conceptualization, Data Curation, Formal analysis, Investigation, Validation, Resources, Writing – Original Draft, Writing – Review & Editing, Visualization, Supervision, Project administration, Funding acquisition

JK: Conceptualization, Data Curation, Investigation, Writing – Review & Editing RM: Conceptualization, Data Curation, Investigation, Writing – Review & Editing

YCL: Conceptualization, Investigation, Resources, Writing – Review & Editing, Supervision, Funding acquisition

MDR: Conceptualization, Data Curation, Investigation, Resources, Writing – Review & Editing, Supervision, Project administration, Funding acquisition

## Methods

### Participants

Thirty-six participants were recruited for the fMRI experiment from the Chicago area (21 female, 14 male, 1 non-binary; mean age 23.42 ± 4.01 with range 18 to 33). No participant was excluded from the analyses. Participants were compensated either monetarily or with the University of Chicago course credits. None of the participants had watched the TV series before the experiment. All participants were right-handed and reported fluency in English. The participants provided informed consent before taking part in the study. The study was approved by the Institutional Review Board of the University of Chicago.

### Stimulus

An episode of the TV series, *This is Us*, season 1 episode 1 (2016, directed by Requa, J. & Ficarra, G. and written by Fogelman, D.), was used as a stimulus in the study (41m 40s). The stimulus was chosen because it is an omnibus, multi-event, multi-character narrative, which has clear event boundaries, dynamic social relationships, and intense emotions that evolve over the course of the episode. Four short narratives play out in the episode in an interleaved order, starring Jack, Kate, Kevin, and Randall as main characters respectively. The four characters’ events are delivered in an independent fashion, except for the connection that all four characters are 36 years old and that Kevin and Kate are siblings. At the end of the episode, there is a big reveal which shows that Jack’s events are from the past and he is the father of Kate, Kevin, and Randall who are triplets.

The episode was segmented into 48 events and scrambled in temporal order. The duration of each segment was on average 52 ± 21s, with a range of 8s to 1m 37s. The events were segmented by the first author based on the director’s cut. We segmented the events if one character’s event changed to another character’s event. Within one character’s continuous event, we segmented when there was a meaningful change in the narrative or temporal/spatial backgrounds. This resulted in 12 events from Jack events, 9 events from Kate events, 9 events from Kevin events, 4 events from Kate & Kevin events, 11 events from Randall events, and the last 3 events that contained the “big reveal”. To scramble these events and group them into blocks in a manner that all blocks hold comparable degrees of importance, eight researchers in the lab (including the first and last authors) rated the importance of the 45 segmented events, excluding the last 3 events which obviously held high importance. The researchers rated how important the events are in the story; if an event provided key information that is critical in comprehending the narrative of the episode, the event would be rated with a high score.

The temporally scrambled 45 events were grouped into 9 blocks such that each block consisted of one event from Jack, Kevin, Kate, and Randall events respectively, plus one additional event that was either Jack event, Randall event, or Kate & Kevin event. The last 3 events were grouped in the original temporal order as one block. The mean block duration was 4m 10s. The event assignments to blocks were done by the first author, so that the summed duration and summed importance of five events per block were comparable across blocks. Five events were ordered in a block such that there were no coincidental regularities in character event sequence (e.g., the likelihood that a Kate event is to be followed by a Kevin event) or the order of event importance (e.g., the likelihood that the blocks are likely to start with an unimportant event and end with the important event). Event assignment and order within each block were fixed.

Having fixed the event order within a block, we scrambled the sequence of nine blocks, to create three scrambled-order groups (**Figure 1A**). We scrambled the order of blocks such that each block tiles the first one-third, middle one-third, and last one-third of the entire block order in the three stimulus groups (e.g., block 1 should be positioned at the beginning for group 1, middle for group 2, and at the end for group 3). Block 10, which contained the last 3 consecutive events of the episode, was always positioned on the 7th run such that the rest of the blocks could be positioned before the “big aha” for the two stimulus groups and after the “big aha” for one stimulus group. By systematically positioning the blocks in sequential order, we were able to compare the effects of contexts, or knowledge gained about the narrative, while holding the perceptual inputs of the event constant for the three stimulus groups.

### FMRI experiment

The fMRI experiment consisted of 10 runs in total. Each run contained three parts: aha button presses during scrambled episode watching (one block of events per run) followed by the verbal explanation of why participants pressed the aha buttons at certain moments, and the verbal response on their personal thoughts about the characters. Participants’ personal thoughts about the characters were not analyzed in the current study.

As participants watched the scrambled episode, they were instructed to pay attention to the video and try their best to speculate what the original story is: the original temporal sequence of the story, causal links that exist between events, and what characters are doing, thinking, and feeling. Participants were asked to press the “aha” button whenever they experienced a subjective feeling of “aha” or a sudden insight. They pressed the button when they grasped what was going on in the story, made sense of the past event that they previously did not understand, realized something new, resolved initial curiosity about any parts of the narrative (even partially), came to have a better understanding of—or even just a speculation about—the general plots, the temporal sequence of the events, causal relationships between events, as well as characters’ behavior, thoughts, feelings, intentions, and relationships with other characters. Whenever they realized that their previous “aha” moments turned out to be incorrect, which is a subjective feeling of “oops”, they also pressed the aha button. All participants received the same instruction from the same instructor prior to the fMRI scans. They were reminded that there were no correct or incorrect answers to the aha button presses, that they should follow their subjective feelings, and that being a frequent or infrequent button presser does not matter in the experiment. The participants were asked to press the button without hesitation when they had even a tiny urge to press. Event onsets were triggered by scan pulses and the timings of onset and offset were recorded.

After the aha button presses during scrambled episode watching was over, participants were presented with screenshots of the events that they pressed the aha button. The screenshots were labeled with numbers in order. Participants were asked to freely explain the reasons why they pressed the aha button at these moments; what they newly understood or learned about the events and characters from watching the events, and how their thoughts about the events and characters changed from watching the events. Participants were given the freedom to either explain their aha experience one by one for each screenshot, or describe their overall aha experience in those moments at an event. Because there were five event segments in each run (with the exception of run 7 which consisted of the last 3 consecutive events of the episode), all aha button presses made at each event were presented together as screenshots. For example, if a participant pressed the button twice in event 2 and once in event 4, they would be given two prompts. One would have two screenshots of event 2, and the next would have one screenshot of event 4. Because the participant did not press the button in events 1, 3, and 5, these prompts were not given. This means task demands for verbal responses differed depending on whether the participant was a frequent or infrequent button presser. Frequent button pressers who pressed the button at every event had to explain multiple screenshots that were consecutively presented for all five events, whereas infrequent button pressors had to explain a few screenshots for less than five events at which they pressed the button. The participants were instructed to talk clearly and accurately with as much detail as possible. If the participants were done talking about the screenshots, they pressed a button to move onto the next prompt. The next prompt started when the next scan pulse was recorded. The timing of the prompt onset was recorded. Only behavioral data were analyzed in this study—we did not analyze neural data acquired during the verbal response phase.

Participants were scanned with a 3T scanner (Magnetom Prisma; Siemens Healthineers, Erlangen, Germany) with a 64-channel head coil. Anatomical images were acquired using a T1-weighted magnetization-prepared rapid gradient echo pulse sequence (repetition time [TR] = 2,300 ms, echo time [TE] = 2.4 ms, field of view = 256 mm × 256 mm, and 1 mm isotropic voxels). Functional images were acquired using a T2*-weighted echo planar imaging (EPI) sequence (TR = 1,200 ms, TE = 25 ms, multiband factor = 4, field of view = 208 mm × 192 mm, and 2 mm isotropic voxels, with 68 slices covering the whole brain).

Visual stimuli were displayed on an LCD 40-inch monitor with a resolution of 1920 × 1080 pixels and a refresh rate of 60 Hz. Auditory stimuli were delivered by Avotec through MRI-compatible comply canal tips earbuds. Participants’ voices were recorded using in-house Matlab software. After the acquisition of raw audio files, our customized software detected the regularity of audio signals coming from scan pulses and denoised it from participants’ voice recordings. Four participants’ voice recordings were not analyzed because the files were not recorded or deprecated during data collection.

Because all participants’ duration for spoken response differed, the experimenters manually stopped the scan when each run ended. Participants’ head motions during scans were monitored using the Framewise Integrated Real-time MRI Monitoring (FIRMM) software. After each run, participants were given appropriate feedback about their motion when needed. The duration of the fMRI experiment varied across participants depending on their amount of verbal responses; it ranged from approximately 1hr 20m to 2hr 15m.

The fMRI study was followed by a post-scan behavioral session comprised of two parts: recounting the scrambled story in its original order and a survey on what they thought about the four main characters. These data were not analyzed in the current study.

### FMRI image preprocessing

The fMRI data were converted to BIDS format. The frames from the movie start time and 3 TRs after the end of the verbal response were extracted from the BIDS output for analyses. All images were deobliqued. Anatomical T1-weighted images were processed with a standard fMRIprep pipeline with surface reconstruction (Esteban et al., 2019). Functional EPI images were also processed with standard fMRIprep pipeline, without slice timing correction, without susceptibility distortion correction, and images registered to MNI152NLin2009cAsym:res-2 template. The EPIs after the fMRIprep were masked with each individual’s MNI-aligned brain mask. We regressed out the following nuisance variables from all voxel time series within the masked brain: 24 head motion parameters, global signal, signals from cerebrospinal fluid and white matter, top-five cCompCor, top-five wCompCor, linear trend, and low-frequency signal with 128 s cut-off. The frames that exceeded a threshold of 0.5 mm FD or 1.5 standardized DVARS, as well as frames that were detected as non-steady-state volumes were censored as NaNs. We applied spatial smoothing using 4 FWHM.

Three participants’ data had more than 10% of total frames censored during video watching due to head motion, thus were excluded from all fMRI analyses (proportion of censored frames: 19.021,16.458, 10.885 %). The remaining 33 participants had minimal head motion during video watching, with an average of 0.665 ± .718 % of the frames censored, ranging from 0% to 2.126%. These participants’ behavioral data were still analyzed.

### Parcellation

We used the subject-specific autogenerated EPI mask and predefined template-registered cortical and subcortical parcels to extract the BOLD time series of voxels. First, using 10 brain masks generated by fMRIprep on the fMRI runs, we created a composite brain mask per participant by taking the intersection. Second, we used the 100 cortical parcels by Schaefer and colleagues (2018) and 16 subcortical parcels by Tian and colleagues (2020). For each of the 116 parcels, BOLD activity time series were extracted from voxels that were contained in both the participant-specific mask and predefined parcels. Note that this process results in some parcels containing different numbers of voxels across participants.

### Data preparation for analysis

During the fMRI experiment, we recorded 1) the onset and offset of the video for each event, and 2) the time participants spent verbally responding to every event in which they pressed the aha button. We limited all our fMRI analyses to moments when participants were watching the events, excluding the verbal response phase. We extracted the BOLD time series from the onset of the first event to the offset of the last event, which was shifted by 4 TRs (4.8 s) to account for the HRF. Each of the extracted voxel time series was z-normalized across time, with censored frames remaining as NaNs.

### Aha button press synchrony

Thirty-six participants’ aha button press moments in seconds were converted to corresponding TRs within each of the 48 segmented events. We recorded button presses up to 2 TRs after each event was over. Dice coefficients were calculated as a measure of button press synchrony. Specifically, 1 TR before and 1 TR after the button press were considered button press moments to compensate for individual differences in response times. Dice coefficients were calculated between every pair of participants, which were grouped into whether the pair belonged to the same scrambled-order group vs. different groups (**Figure 1D**).

### Intersubject correlation (ISC) analysis

The BOLD time series of all voxels within each of the 100 cortical and 16 subcortical parcels were averaged to represent the mean activity time series of a parcel. Pearson’s correlations of the mean activity time series were calculated across all participant pairs. Participant pairs were grouped based on whether the pair belonged to the same scrambled-order group or different groups. The ISC difference was calculated as the mean of same-group ISC minus the mean of different-group ISC (calculated after Fisher’s *r*-to-*z* transformation). The ISC difference was calculated for every parcel and visualized in the surface brain using nilearn.plotting.plot_surf_stat_map (**Figure 1E**).

### Coding of verbal responses and causal relationship

Thirty-two participants’ verbal explanations of their aha button presses (recorded inside the fMRI) were transcribed verbatim. After listening to the explanations, we defined eight categories that encompassed a broad spectrum of participants’ verbal responses (**Supplementary Table S1**). Three of the authors categorized the explanations associated with each button press into one or more of these categories. If a participant explained the screenshots separately, different categories were assigned to the corresponding button presses, whereas if a participant explained multiple screenshots together with a single explanation, the same categories were assigned to the corresponding button presses.

For the “memory retrieval” category, which was labeled when there were either explicit or implicit mentions of past events, the three raters annotated which event(s) were mentioned. This allowed us to create a memory retrieval matrix for each participant by marking +1 if a past event was mentioned at the insight moment of a current event. This directed matrix was then unscrambled in temporal order and averaged across all 32 fMRI participants to create one undirected memory retrieval matrix. We did not include run 7 in the analyses, as the majority of explanations were “big aha” when participants understood the family relationship between the four characters.

Causal relationships of event pairs were mapped by the same three raters, on a scale of 0 (no causal relationship) to 4 (very high causal relationship). The instruction and definition of the causal relationship provided to the raters are shown in **Supplementary Table S2** (Trabasso et al., 1989; Antony et al., 2024).

### Narrative feature extraction

Five narrative features—semantics, characters, places, low-level visual, and low-level audio features— were extracted to represent each of the 48 events (**Supplementary Figure S1A**). To represent semantics, the contents of each event were annotated in detail by the first author. The annotations in descriptive sentences, together with the transcripts of dialogues within each event, were fed into the pre-trained GPT-2 language model (transformers.GPT2Model) (Radford et al., 2019). The 768-dimensional embeddings of the last hidden layer were extracted from every meaningful word (transformers.AutoTokenizer.from_pretrained(“gpt”)) and averaged across words within each event. This resulted in a 768-dimensional embedding vector respective to 48 events. Which characters appeared in each event and where the event took place were coded as one-hot vectors by the first author. Character vectors represented the appearance of 4 main characters, Jack, Randall, Kate, and Kevin, and 9 sub-characters. Place vectors included a total of 15 unique indoor (e.g., Jack’s bedroom, Randall’s office, television set, hospital, support group, restaurant) and outdoor (e.g., soccer field, in front of Kate’s house) settings. The low-level visual features were characterized by four features—hue, saturation, pixel intensity, and motion energy—which were estimated per frame and averaged across frames within an event. Hue, saturation, and pixel intensity were estimated with MATLAB’s rgb2hsv function. Motion energy was estimated with pliers.extractors.FarnebackOpticalFlowExtractor function. The low-level audio features were represented with four features—amplitude and pitch of left and right stereos—which were estimated per frame and averaged across frames within an event. Amplitude and pitch were estimated using audioread and pitch functions in MATLAB. The event-by-event narrative feature similarity matrix was created by taking Pearson’s correlation coefficient, which was Fisher’s *r*-to-*z* transformed for further analyses.

Lastly, to represent how close in time the event pairs were in the narrative’s original sequence, events were sorted in their original order respectively for the four characters’ narratives. The events where Kevin and Kate appeared together were included in both Kate’s and Kevin’s narratives. If an event pair was continuous in time, the highest score (score 14) was given, and if an event pair was furthest in time (i.e., the first and last event within a character narrative that was separated by 13 events in between) the lowest score (score 1) was given. A score of 0 was given to event pairs of different character narratives (**Supplementary Figure S1A**).

### Neural pattern shift detection

The hidden Markov model (HMM)-based segmentation model was implemented using the function brainiak.eventseg.event.EventSegment. We applied the HMM on the multivariate BOLD activity of all voxels corresponding to each of the 100 cortical parcels, for each of the 48 events, and for each of the 33 participants, independently. We intentionally did not apply the model to subcortical parcels because subcortical voxel activity shifts rapidly at every TR, which does not meet the HMM’s assumption that a latent event prolongs for a continuous time period. NaN values due to censoring were linearly interpolated from the neighboring frames. For each event’s time series, we iterated over a different number of latent states (i.e., K) for optimal hyperparameter selection. If the event duration was smaller than 10 TRs, we did not apply the HMM and considered the event as a continuous time series. If the event duration was less than 20 TRs, we started event iterations from K = 2, whereas we started from K = 3 for events with durations over 20 TRs. For the maximum number of K in the hyperparameter search, we took the floor of each event duration divided by 3, such that the average duration of each event is at least 3 TRs. We applied the HMM across iterations over possible Ks, and calculated the difference in voxel pattern similarity within the same event compared to different events. In this comparison, we controlled for the separation in time when computing the voxel pattern similarity between different time points. That is, if the within-event similarity was calculated between TRs separated in 4 time steps, the across-event event similarity was calculated between TRs also separated in 4 time steps. We chose K which exhibited the largest within-event similarity compared to across-event similarity. However, if an event showed a monotonic decrease or increase in within-compared to across-event similarities across a range of K, we considered it a poor model fit and did not apply the HMM for those events (i.e., the entire event was considered as a single event). The “boundaries” detected by the HMM represented moments when multivariate voxel representation patterns shifted from one time segment to the next. These moments of neural pattern shifts were annotated with index 1, whereas other moments were indexed with 0.

To align these moments of neural pattern shifts with the aha button press moments, we took the boundary indices (1 for moments when the pattern shifted, 0 for the rest) at -5 to 3 TRs from all aha button presses for each participant and averaged them across button press moments. Time frames that did not exist within this range (e.g., aha button press that was made less than 5 TRs since the onset of the event) were treated as NaNs. For a statistical test, we iterated the same analysis 10,000 times with the aha button press moments randomly shuffled in time. The likelihood of the neural pattern shifts averaged across all 33 participants was compared to the chance distribution at each time step, which was corrected for multiple comparisons across parcels and time steps.

### Neural reinstatement of causally related past events

Twenty-nine participants’ data were analyzed; 3 excluded for head motion and 4 excluded for the failure of behavioral response recording. To estimate the neural reinstatement effect, for each aha button press, we compiled neural patterns of all past events that occurred in prior fMRI runs. These patterns were multiplied by their degrees of causal relationship to the current event which were taken from the causal relationship matrix. The weighted linear sum of the past events represented the integrated pattern of the causally related past events. Pearson’s correlation between the integrated past pattern and the current pattern was calculated ranging from -10 to 3 TRs from the HRF-shifted aha button presses. Our analyses were restricted to 19 cortical regions that significantly shifted neural patterns at 2 TRs prior to aha button presses. For the same time frame (from -10 to 3 TRs) and every button press, we extracted a binary measure of whether the neural pattern shift was detected by the HMM at -2 TR. For the same aha button press, we also extracted a binary measure of whether past event(s) were mentioned in the participant’s verbal explanation of their insight. If any of the three coders judged that the insight accompanied retrieval of past event(s), we considered it memory retrieval.

The aha button presses were categorized based on binary measures of neural pattern shifts at -2 TRs and behavioral memory retrieval. Based on the observation of **Figure 4A**, we considered that neural reinstatement occurs at approximately -7 to -3 TRs from aha button presses. Neural reinstatement effects in this time frame were averaged (calculated after Fisher’s *r*-to-*z* transformation).

Logistic generalized linear mixed effect models were conducted using the lme4 library in R, since the neural shift and behavioral retrieval measures were both binary (“binomial” family in glmer function): neural pattern shift ∼ neural reinstatement + 19 regions + (1|subject), and behavioral retrieval ∼ neural pattern shift + neural reinstatement + neural pattern shift × neural reinstatement + 19 regions + (1|subject). With regard to the latter model, we also ran a model where neural pattern shifts detected on either 2 TR or 1 TR prior to aha button presses were indicated as 1. Chi-squared tests were conducted between each of these models and the reduced models as we removed each one of the main effects or the interaction effect individually.

### Event-by-event neural pattern similarity and causal relationship

For each of the 100 cortical and 16 subcortical parcels, we averaged multivariate voxel activity pattern across time for each of the 48 events across 10 runs. Taking the correlation coefficients of (number of voxels) × (48 events in a scrambled order) resulted in a (48 events) × (48 events) BOLD pattern similarity matrix for each participant, which was Fisher’s *r*-to-*z* transformed (**Figure 5A**). This was correlated with the causal relationship matrix in the scrambled order using Spearman’s rank correlation. The average Spearman’s *rho* value across 33 participants was compared with a chance distribution where *rho* was calculated between the shuffled BOLD pattern similarity matrix and the causal relationship matrix (10,000 iterations). This was iterated across 116 parcels, with the significance corrected for 116 comparisons (FDR-*p* < 0.05).

In the same manner, we calculated partial rank correlation (pingouin.partial_corr), where we correlated BOLD pattern similarity with causal relationship while controlling for 6 narrative feature matrices. To avoid collinearity, 6 narrative feature matrices were converted into 6 principal components prior to calculation.

## Supplementary Information

### Supplementary Figures

**Supplementary Figure S1.**
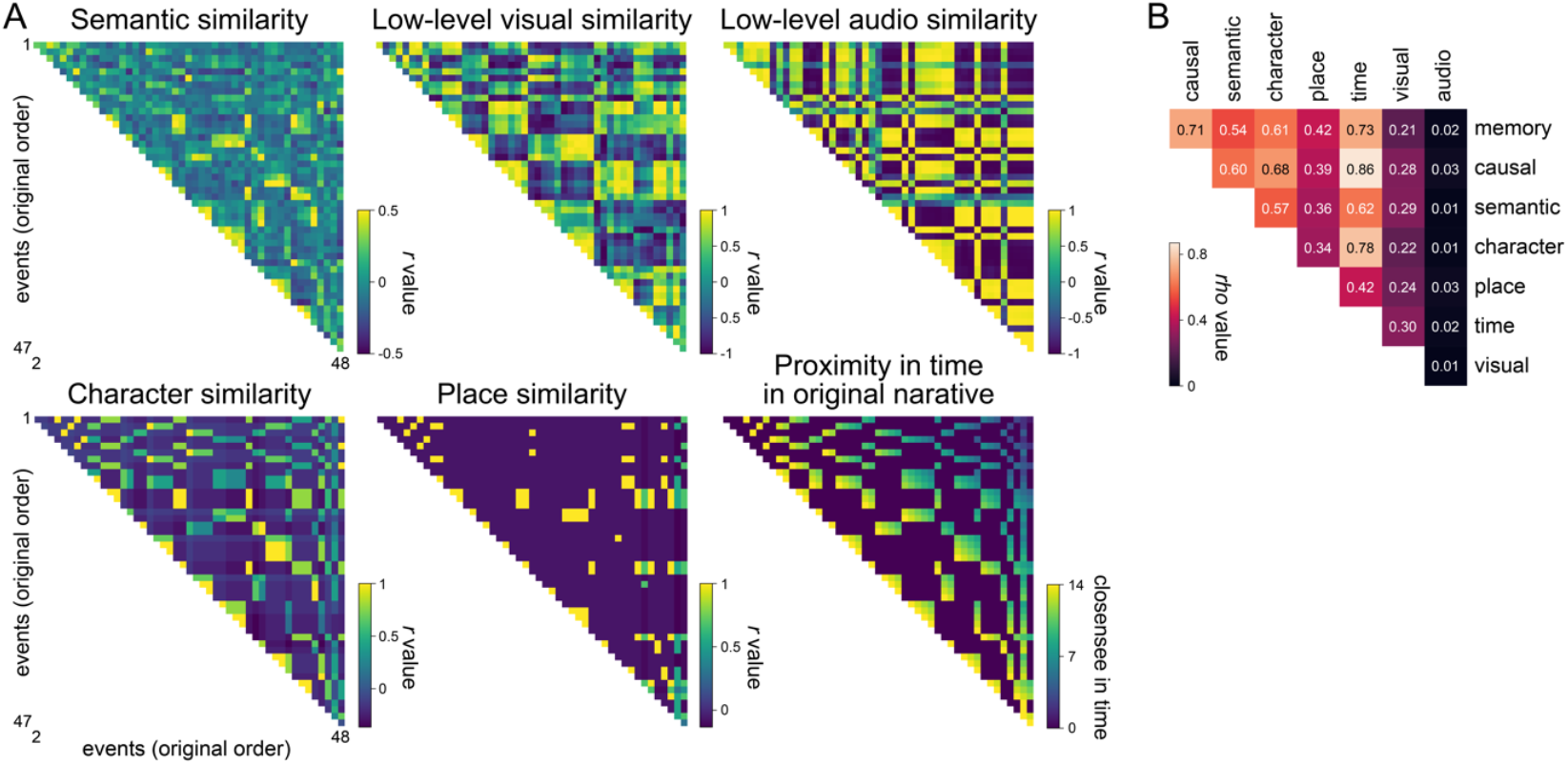
Event-by-event narrative feature matrices and their relationship to one another. **(A)** Narrative feature matrices in the original temporal sequence. The semantic similarity matrix denotes similarities in semantic contents between pairs of events. The semantics of each event were represented from the annotation of event contents where all meaningful words were transformed into 768 dimensional embeddings by a pre-trained GPT-2 language model and averaged per event. Each event was binarily coded as to what characters appeared (13 dimensional one-hot vectors: 4 main characters and 9 sub-characters) and where the event took place (15 dimensional one-hot vectors of unique indoor and outdoor settings). The low-level visual features (hue, saturation, pixel intensity, and motion energy) and auditory features (amplitude and pitch) were extracted from every frame of the video and audio, which were averaged to represent the summary features of each event. Pearson’s correlation coefficients were calculated for these narrative features for every pair of events. For the time proximity matrix, events were primarily grouped into independent 4 characters’ narratives. Event pairs nearby in time were rated with high value and far in time were rated with low value. Unrelated event pairs were coded with 0. **(B)** Pairwise relationship between narrative features. Spearman’s rank correlation was calculated between every pair of narrative feature matrices, including memory retrieval and causal relationship matrices.

**Supplementary Figure S2.**
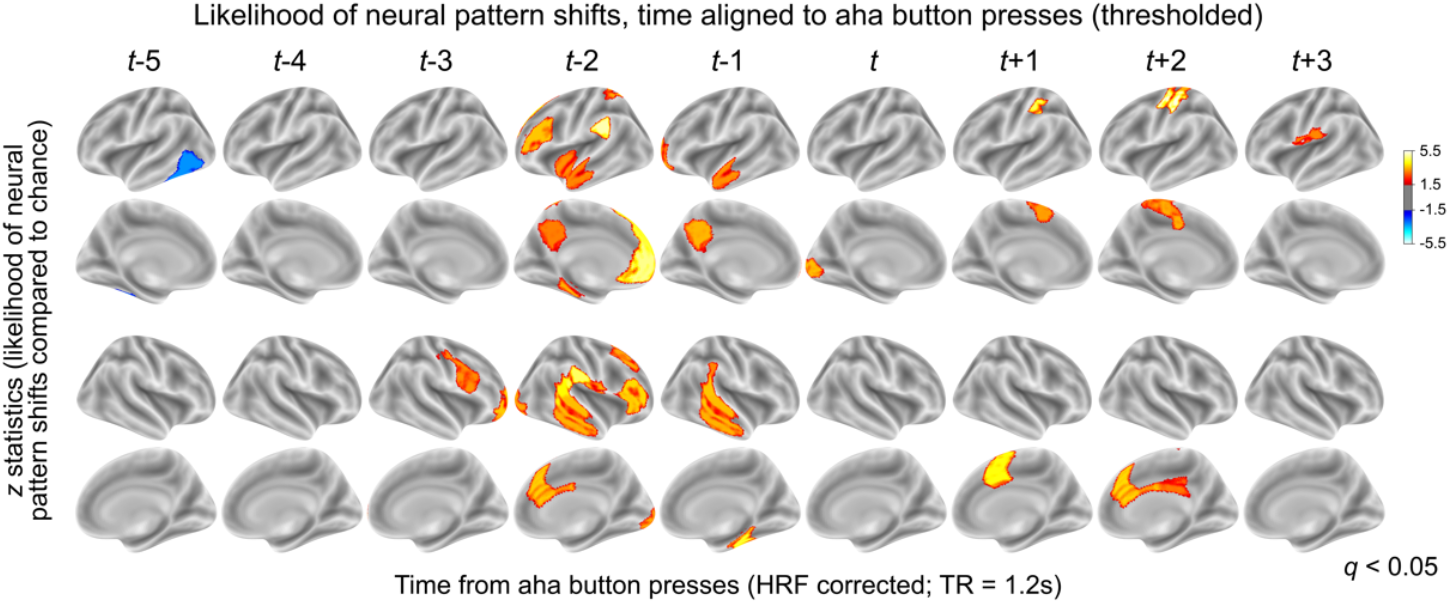
Thresholded *z* statistics of **Figure 3C**. Likelihood of the HMM-detected neural pattern shifts compared to a chance distribution with randomly shuffled aha responses. Colors indicate z statistics computed in comparison to the chance which was generated at each parcel and each time point, thresholded at FDR-*p* < 0.05.

### Supplementary Tables

**Supplementary Table S1.** Thirty-two participants’ explanations of their aha button presses coded into different categories. Categories were not determined *a priori* but were generated by the first author during coding, based on participants’ verbal responses. The verbal response for each aha button press was able to belong to multiple categories. The proportion of button presses belonging to each category compared to the total number of button presses was calculated, and averaged across three raters and participants to derive the probability (%) estimate. Reliability was calculated as average Spearman’s correlations between all pairs of raters’ category probability estimates of 32 participants.

| Category | Description | Average (SD) % | Reliability ($\rho$) |
| --- | --- | --- | --- |
| Character | Comprehension of the characters' traits, behavior, thoughts, or feelings. | 53.76 (13.97) | 0.649 |
| Current event understanding | Comprehension or description of what's happening in the current event. | 47.49 (15.24) | 0.483 |
| Memory retrieval | Explicit or implicit mention of past events. | 41.15 (16.64) | 0.762 |
| Character relationship | Comprehension of the relationship between characters. | 23.00 (7.41) | 0.541 |
| Temporal order | Comprehension of the original temporal order of the event(s). | 17.59 (13.42) | 0.787 |
| Inference | Inference on narrative contents or connecting links between events. Prediction of future events. | 12.63 (6.7) | 0.393 |
| Causal understanding | Comprehension of causal structure of the narrative, or causal relationship between events. | 11.64 (6.75) | 0.315 |
| Oops | Realizing prior comprehension had been incorrect and/or correcting for it. | 6.68 (5.19) | 0.444 |

**Supplementary Table S2.**
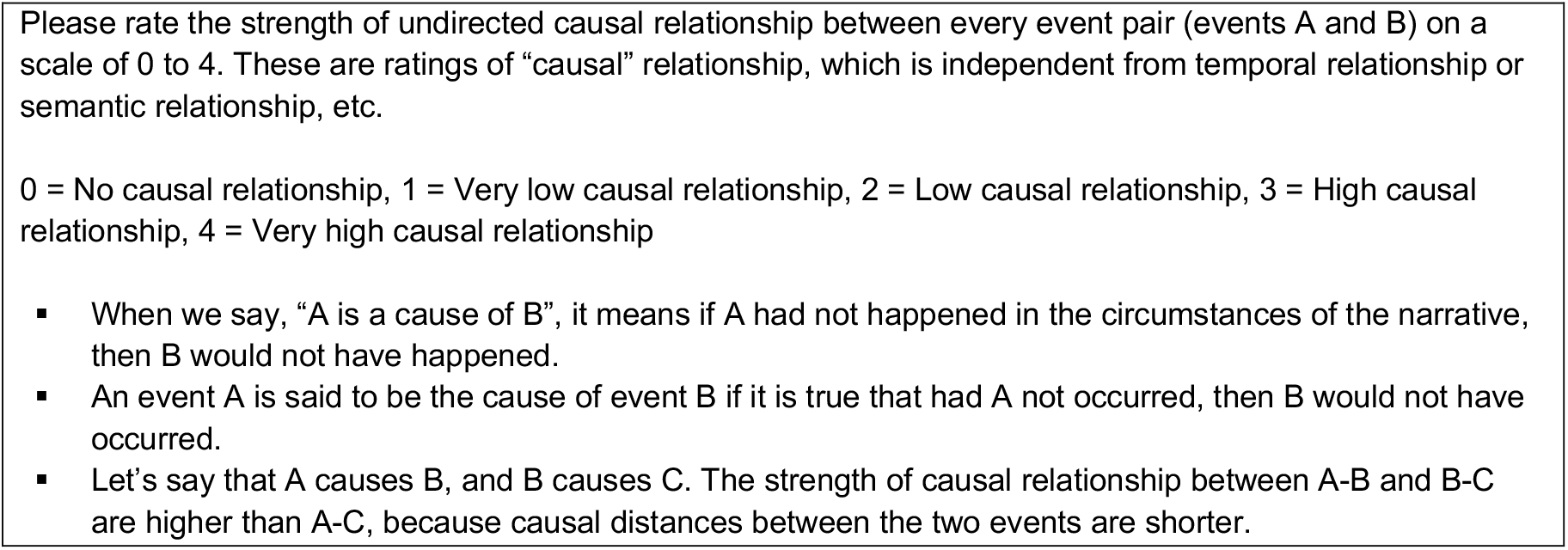
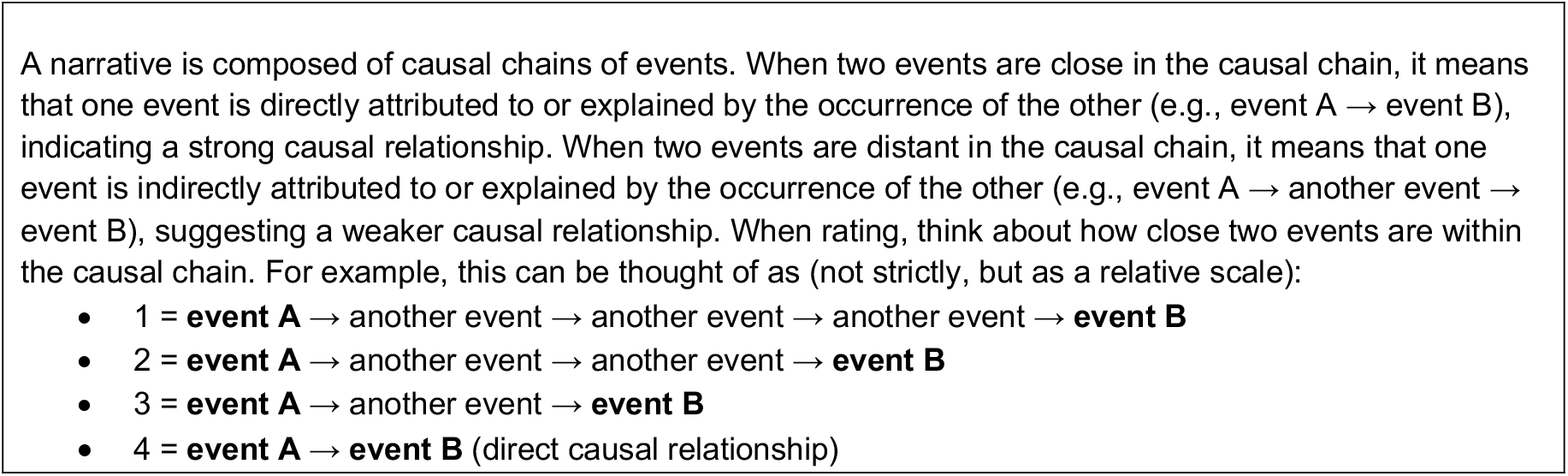
Definition of causal relationship and the instruction used in causal relationship coding.

**Supplementary Table S3.** Nineteen cortical areas that exhibited significant neural pattern shifts at 2 TRs prior to aha button presses. Region names and numbers come from Schaefer et al.’s (Schaefer et al., 2018) MNI templated-aligned 100 cortical parcellation scheme.

|  |  |  |  |
| --- | --- | --- | --- |
| 19 | 17Networks_LH_DorsAttnB_PostC_3 | 52 | 17Networks_RH_VisCent_ExStr_2 |
| 21 | 17Networks_LH_SalVentAttnA_ParOper_1 | 63 | 17Networks_RH_SomMotB_S2_2 |
| 22 | 17Networks_LH_SalVentAttnA_Ins_1 | 71 | 17Networks_RH_SalVentAttnA_ParOper_1 |
| 32 | 17Networks_LH_ContA_PFCI_1 | 77 | 17Networks_RH_SalVentAttnB_PFCmp_1 |
| 39 | 17Networks_LH_DefaultA_pCunPCC_1 | 81 | 17Networks_RH_ContA_PFCI_1 |
| 40 | 17Networks_LH_DefaultA_PFCm_1 | 83 | 17Networks_RH_ContB_Temp_1 |
| 41 | 17Networks_LH_DefaultB_Temp_1 | 90 | 17Networks_RH_DefaultA_PFCd_1 |
| 44 | 17Networks_LH_DefaultB_PFCd_1 | 95 | 17Networks_RH_DefaultB_PFCv_2 |
| 49 | 17Networks_LH_DefaultC_PHC_1 | 99 | 17Networks_RH_TempPar_2 |
|  |  | 100 | 17Networks_RH_TempPar_3 |

### Supplementary Text

Baldassano et al. (2017) used the hidden Markov model to show that moments of voxel pattern shifts correspond to moments of perceived event boundaries. Because we used the same method to show that voxel pattern shifts correspond to moments of insight, we asked about the relationship between moments of insight and perceived event boundaries.

A behavioral experiment was conducted using the same TV episode stimulus in three versions of scrambled orders (N=36 of a different participant sample, N=12 in each scrambled-order group). Participants were instructed to press a button when a meaningful unit ended and another began, which is the replication of event boundary button press instruction given by prior studies (Newtson, 1973; Zacks et al., 2010; Ben-Yakov and Henson, 2018; Michelmann et al., 2021; Bezdek et al., 2023).

Participants were recruited from the University of Chicago and compensated either monetarily or with course credits (21 female, 16 male; mean age 20.11 ± 2.157 with range 18 to 29). We did not screen volunteers based on whether they watched the TV series before the experiment, but we excluded volunteers who participated in our fMRI experiment. The participants provided informed consent before taking part in the study. The study was approved by the Institutional Review Board of the University of Chicago.

Participants pressed the button at similar moments in time (*z* = 86.312, *p* < 0.001, compared to a chance distribution where the button press moments were randomly shuffled 10,000 times), which was significantly higher than the synchrony of the aha button responses (Wilcoxon rank-sum test: *z* = 18.015, *p* = 1.49e-72). In contrast to the aha button press experiment, the synchrony in perceived event boundaries did not differ depending on whether the participant pairs belonged to the same or different scrambled-order groups (Wilcoxon rank-sum test: *z* = 0.682, *p* = .495). This means perceived event boundaries did not differ depending on differences in knowledge or context about the narratives.

Importantly, we calculated dice coefficients between 36 participants’ aha button presses and 36 participants’ event boundary button presses for all pairs that belonged to the same scrambled-order group. The synchrony between aha and event boundary button presses was significantly above chance, where button presses were randomly shuffled (*z* = 20.248, *p* < .001). This suggests that the likelihood of insight moments overlapping with perceived event boundaries is statistically higher than what is expected by chance.

**Supplementary Text S1**. Relationship between moments of perceived event boundaries and moments of insight.

